# Naturally secreted bacterial outer membrane vesicles: potential platform for a vaccine against *Campylobacter jejuni*

**DOI:** 10.1101/2020.10.16.342261

**Authors:** Ankita Singh, Afruja Khan, Tamal Ghosh, Samiran Mondal, Amirul Islam Mallick

**Author notes:** Corresponding Author *Corresponding Author’s Address*: Dr. Amirul Islam Mallick, Associate Professor, Department of Biological Sciences, Indian Institute of Science Education and Research Kolkata, Mohanpur, Nadia, West Bengal, 741246, India. Phone: +**91 033 2502 8000** (Ext 1221).

## Abstract

Acute diarrheal illness and gastroenteritis caused by *Campylobacter jejuni* (*C. jejuni*) infection remain significant public health risks in developing countries with substantial mortality and morbidity in humans, particularly in children under the age of five. Despite improved global awareness in sanitation and hygiene practices, including food safety measures, *C. jejuni* infections continue to rise even across the developed nations and no vaccine is currently available for humans. Genetic diversities among *C. jejuni* strains as well as limited understanding of immunological correlates of host protection remain primary impediments for developing an effective vaccine against *C. jejuni*. Given the role of bacterial outer membrane-associated proteins in intestinal adherence and invasion as well as modulating dynamic interplay between host and pathogens, bacterial outer membrane vesicles (OMVs) have emerged as potential vaccine platforms against a number of enteric pathogens, including *C. jejuni*. In the present study, we describe a mucosal vaccine strategy using chitosan (CS) coated OMVs (CS-OMVs) to induce specific immune responses against *C. jejuni* in mice. However, considering the challenges of mucosal delivery of OMVs in terms of exposure to variable pH, risk of enzymatic degradation, rapid gut transit, and low permeability across the intestinal epithelium, we preferentially used CS as a non-toxic, mucoadhesive polymer to coat OMVs. Mucosal administration of CS-OMVs induced high titre of systemic (IgG) and local (secretory IgA) antibodies in mice. The neutralizing ability of secretory IgA (sIgA) produced in the intestine was confirmed by *in vitro* inhibition of cell adherence and invasion of *C. jejuni* while *in vivo* challenge study in OMVs immunized mice showed a significant reduction in cecal colonization of *C. jejuni*. Moreover, to investigate the immunological correlates of the observed protection, present data suggest OMVs driven T cell proliferation with an increased population of CD4^+^ and CD8^+^ T cells. In addition to antibody isotype profile, significant upregulation of IFN-γ and IL-6 gene expression in mesenteric lymph nodes collected from OMVs immunized mice further suggests that mucosal delivery of OMVs promotes a Th1/Th2 mixed type immune responses. Together, we provide strong experimental evidence that as an acellular and non-replicating canonical end product of bacterial secretion, mucosal delivery of OMVs may represent a promising platform for developing an effective vaccine against *C. jejuni*.

**Author Summary:** Despite the loss of 7.5 million disability-adjusted life years, which is over and above any other globally prevalent enteric or enterotoxigenic pathogens, *C. jejuni* remains a neglected foodborne pathogen, particularly in tropical countries. Even with the improved global awareness in sanitation and hygiene practices, including food safety measures *C. jejuni* infections continue to rise globally and no vaccine is currently available for humans. In light of the importance of the diverse cargo selection by bacterial OMVs, the present study describes a mucosal vaccine strategy using chitosan-coated OMVs to induce specific immune responses against *C. jejuni* in mice. We provide here strong experimental evidence that as a non-replicating canonical end product of bacterial secretion, mucosal delivery of OMVs represents an attractive vaccine platform against *C. jejuni*.

## Introduction

*Campylobacter jejuni* (*C. jejuni*) is a common etiological agent associated with an acute, self-limited gastrointestinal illness characterized by diarrhea, fever with several extraintestinal complications such as Guillain-Barre Syndrome (GBS), Reactive Arthritis (RA), Inflammatory Bowel Disease (IBD) [1–3]. Despite the concerted effort over the past two decades towards developing an effective strategy to control *C. jejuni* transmission to humans, except for biosecurity measures, no vaccine is currently available [4,5]. Moreover, the alarming trends in the rapid emergence of antibiotic resistance among *C. jejuni* have essentially entailed the need for innovative approaches towards developing an effective vaccine platform against *C. jejuni* [6–8]. Such a platform should base on a clear rationale for choosing the immunological and microbial biomarkers that are directly involved in host-pathogen interaction. To this end, bacterial outer membrane vehicles (OMVs) are known to carry diverse cargoes, including proteins that are actively associated with bacterial adhesion and invasion to participate in dynamic interplay at the host-pathogen interface [9–13].

As a generalized constitutive secretion system, in addition to the outer membrane and periplasmic contents, OMVs often carry nucleic acids, toxins, virulence factors as intrinsic secretory components [9, 14–20]. To venture the function of OMVs in bacterial infections and communication, recent proteomic analysis of *C. jejuni* OMVs have identified more than 150 proteins, including periplasmic, outer membrane-associated, inner-membrane as well as cytoplasmic proteins [9–12]. Importantly, protected within a lipid bilayer, OMVs content often survives longer in harsh extracellular environments than the free form of macromolecules released through other secretory mechanisms. Therefore, as an acellular and non-replicating canonical end product of bacterial secretion, the use of naturally secreted OMVs could be a step forward to a potential vaccine platform against many gut pathogens including *C. jejuni* [9, 21–23].

Because as a mucosal pathogen, *C. jejuni* primarily adhere and replicate in the intestinal epithelial cells and a strong local immune response at the mucosal surface is crucial for effective control of *C. jejuni*, we specifically focused our efforts to develop a mucosally deliverable immunization modality in a murine model [24]. However, vaccines targeting mucosal surfaces are often challenging because of the risk of pH susceptibility, enzymatic degradation, rapid gut transit, and low permeability across the intestinal epithelium [24–27]. To surmount these limitations, in the present study, we chose to use chitosan (CS) as a protective shield owing to the large surface area, excellent mucoadhesive property, biodegradability, low immunogenicity with enhanced ability to adsorb on Microfold cells (M-cells) [28–38].

In this study, we demonstrated that mice mucosally administered with CS coated OMVs (CS-OMVs) induced a high titre of systemic (IgG) and local (sIgA) antibodies. The neutralizing ability of the sIgA produced in the intestine was confirmed by *in vitro* inhibition of cell adherence and invasion of *C. jejuni* while *in vivo* challenge study in immunized mice showed a significant reduction in cecal load of *C. jejuni*. Further, to investigate the immunological correlates of the observed protection, present data suggest OMVs driven T cell proliferation with increased CD4^+^ and CD8^+^ T cells population. In addition to antibody isotype profile, transcriptional upregulation of IFN-γ and IL-6 genes in mesenteric lymph nodes collected from immunized mice further indicates that mucosal administration of OMVs could promote balanced type Th1/Th2 immune responses.

Collectively, data presented herein suggest that with convenience and ease of processing, naturally secreted OMVs may constitute a promising platform towards developing an effective acellular mucosal vaccine against *C. jejuni* for humans.

## Results

### Spherical morphology of naturally secreted OMVs of *C. jejuni*

Electron microscopic analysis of OMVs released from *C. jejuni* suggests spherical morphology of the vesicles with an approximate diameter of ~110 nm (SEM) and ~130 nm (TEM). The absence of bacterial debris in micrographs confirmed the purity of OMVs fractions (Fig 2A and 2B). In terms of total protein content, it was estimated to be 100 μg in 200 mL of a fresh culture of *C. jejuni*.

### Biophysical characterizations of OMVs suggest a change in size and net surface charge with enhanced *in vitro* stability when coated with CS

The mean hydrodynamic diameter of OMVs and CS-OMVs was measured to be ~150 nm and ~220 nm, respectively. In addition, CS coating of OMVs reduced the net surface charges from –23.4 mV to –8.2 mV and enhanced the *in vitro* stability in terms of size distribution, uniformity, and PDI index at different physiological conditions (S1 Fig, Table S1). Similar observations with respect to the overall size of CS-OMVs were recorded by FESEM and TEM analysis, which were ~157 nm and ~165 nm, respectively (Fig 2A and 2B, Table 2).

### Co-incubation of *C. jejuni* with OMVs enhanced bacterial invasion of host cells

To determine whether the interaction of OMVs with target cells have an influence on the invasion ability of *C. jejuni*, gentamicin protection assay was performed. Human INT407 cells infected with *C. jejuni* in the presence of exogenous OMVs showed an enhanced capacity of *C. jejuni* invasion in a dose-dependent manner (5 μg/mL, 10 μg/mL, 20 μg/mL) (*C. jejuni* + OMVs Vs *C. jejuni* only, *P* ≤ 0.01) (Fig 1).

**Fig 1.**
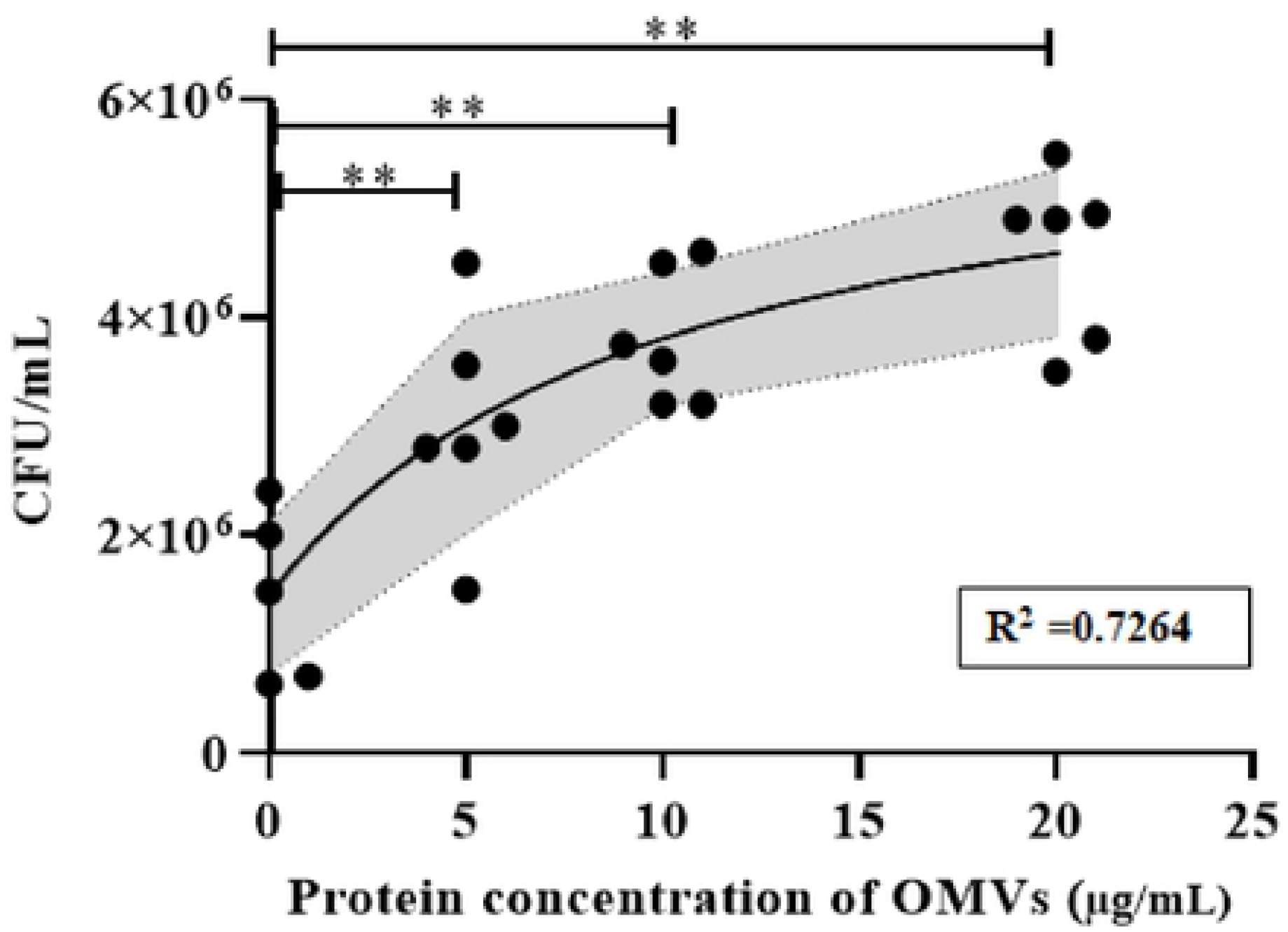
Effect of OMVs on *C. jejuni* invasion of human INT407 cells. The confluent monolayer of human INT407 cells was co-incubated with different concentrations of OMVs (5 μg/mL, 10 μg/mL, and 20 μg/mL) and *C. jejuni* (MOI 1:100) for 3 h at 37 °C in the presence of 5 % CO_2_. After 3 h, cells were washed and treated with gentamicin (150 μg/mL) to kill extracellular and adhered bacteria, followed by incubation for an additional 2 h. Post incubation, infected cells were lysed with 1 % Triton-X 100 to recover the invaded bacteria present intracellularly. Data indicate a significant increase in *C. jejuni* invasion of host cells in the presence of OMVs when compared to *C. jejuni* alone. The assay was performed in triplicate, and regression value (R^2^ = 0.74) was calculated through a non-linear regression curve using GraphPad Prism software (Version 8). Data represent Mean CFU/mL ± SE of two independent experiments. Asterisks indicate a statistically significant difference (\*\**P* ≤ 0.01) with respect to *C. jejuni* alone (without co-incubation with OMVs).

**Fig 2.**
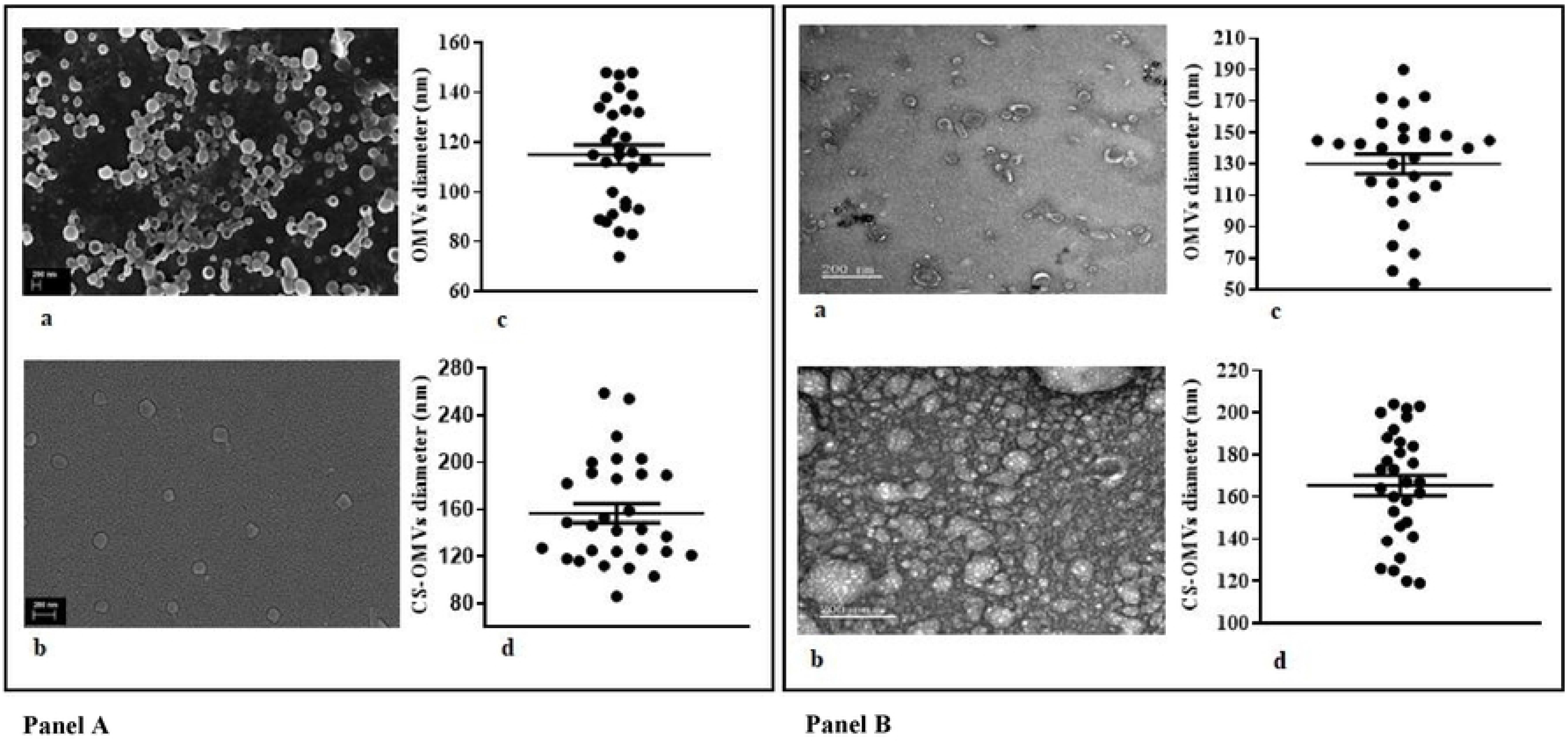
Morphological features and size distribution of OMVs and CS-OMVs by FESEM and TEM analysis. **(A)** Scanning Electron micrograph of OMVs (a) and CS-OMVs (b) shows the spherical shape of the isolated OMVs. The statistical analysis of vesicle size distribution using image J software shows an average size of ~110 nm (c) and ~157 nm (d) for the free OMVs and CS coated OMVs, respectively. **(B)** Transmission Electron micrograph of OMVs (a) and CS-OMVs (b) shows spherical shaped vesicles with the average size for OMVs ~ 130 nm (a) whereas for CS-OMVs ~165 nm (b). The scale bar is 200 nm.

### Mucosal administration of CS coated OMVs induced strong local (sIgA) immune responses in mice

To assess the mucosal immune responses imparted by OMVs administration in mice, faecal pellets and intestinal lavages were collected at day 7 post last immunization and processed by indirect ELISA. The comparative analysis of mean antibody titre of local sIgA present in faecal soup and lavages indicates a substantial increase in sIgA titre in mice that received either CS-OMVs or IFA-OMVs compared to the control mice (received PBS only) (*P* ≤ 0.01) (Fig 3B and 3C).

**Fig 3.**
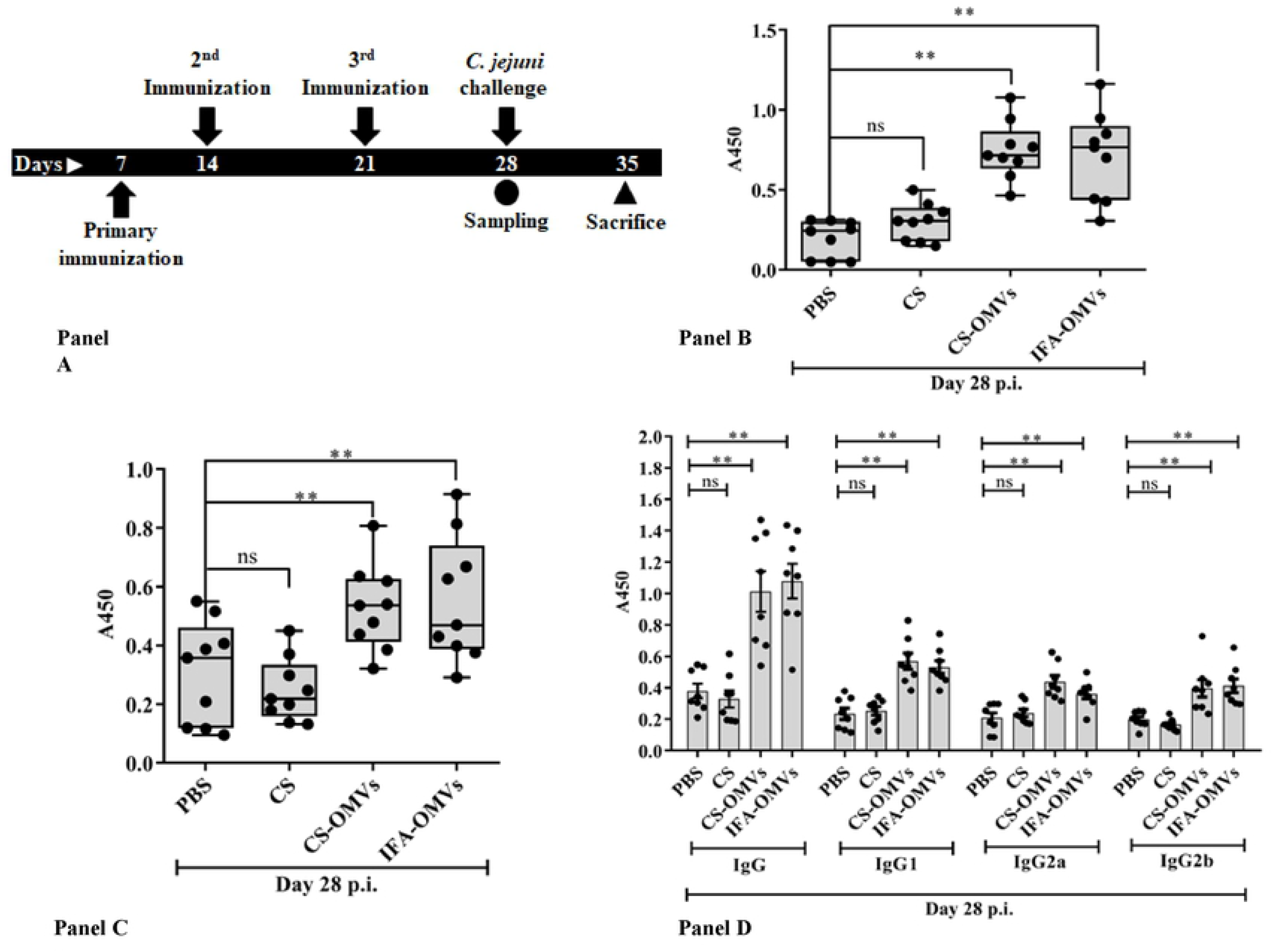
Mice immunization schedule and antibody responses. **(A)** Schematic representation of the mice immunization regimen used in this study. The experimental mice were immunized at indicated time intervals. Circle (●) and triangle (▲) indicate the time points of mice sampling and sacrifice, respectively. At day 28 post first immunization (p.i.), half of the mice were sacrificed to obtain blood, intestinal lavages, faecal pellets, spleen, and mLN samples, whereas on day 35 p.i. the remaining mice were sacrificed to collect cecum. **(B-D)** Comparison of OMVs specific mucosal (sIgA) and systemic (IgG) immune responses in the samples collected at day 7 post last immunization. **(B)** The mean sIgA antibody titre in intestinal lavages (1:8 dilution) and **(C)** faecal soup (1:16 dilution) obtained from the mice of each experimental group showed a substantial increment in sIgA levels in mice either immunized orally with CS-OMVs or injected (s/c) with IFA-OMVs compared to the control mice those received PBS only. **(D)** Comparative analysis of serum IgG isotypic profile based on IgG1, IgG2a, and IgG2b subclasses among different groups showed a significantly higher systemic antibody response and indicated a balanced Th1/Th2 profile (Immunized Vs Controls). Each bar represents the mean antibody titre in the sera samples (1:40 dilution) collected from different experimental groups. Data represent Mean absorbance (A450) ± SE of two independent experiments. Asterisks indicate a statistically significant difference (\*\**P* ≤ 0.01) with respect to the PBS control group.

### CS-OMVs mediated induction of mixed type Th1/Th2 immune responses in mice

Systemic antibody responses in the sera of immunized mice were further examined by assessing the presence of OMVs specific serum IgG level as well as different subclasses (IgG1, IgG2a, and IgG2b) by indirect ELISA. With respect to total IgG responses, animals that received either CS-OMVs or IFA-OMVs showed a significant rise in antibody titre at day 7 post last immunization (*P* ≤ 0.01). Critical analysis of IgG subclasses suggests substantial increment in IgG1, IgG2a, and IgG2b in the sera of CS-OMVs immunized mice followed by IFA-OMVs injected mice (*P* ≤ 0.01) (Fig 3D).

### OMVs driven cellular immune responses in immunized mice

To determine the mucosal delivery of OMVs in inducing specific cellular responses, *in vitro* splenocyte proliferation assay was performed at day 7 post last immunization. Significant increase in cell proliferation rate as revealed by higher stimulation index in mice that received either CS-OMVs (S.I=4.6) or IFA-OMVs (S.I=4.8) (*P* ≤ 0.01) indicate effective priming of T cells by OMVs. In contrast, no detectable response was found in the case of control groups (PBS or CS administered mice) (Fig 4A).

**Fig 4.**
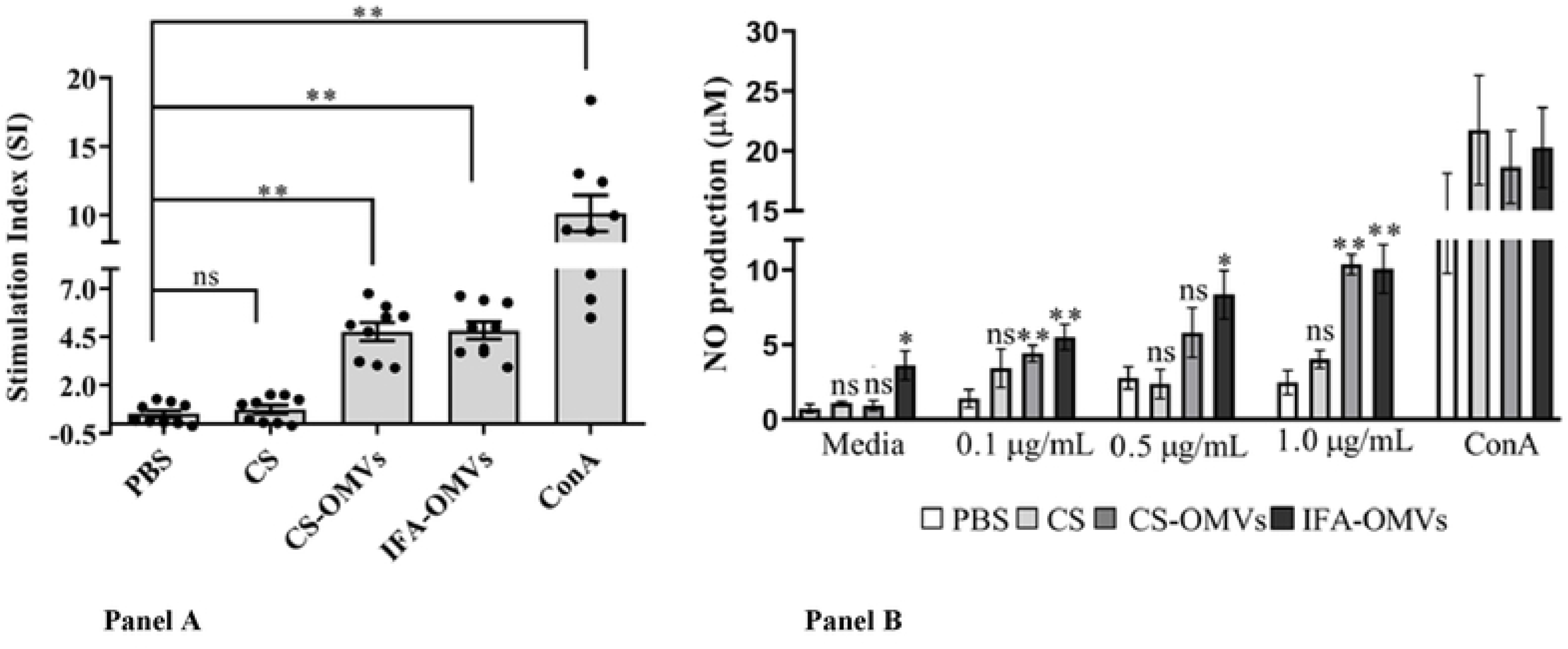
*In vitro* splenocyte (lymphocyte) proliferation and NO production. (**A**) OMVs specific cell-mediated immune responses were measured by BrdU incorporation into nucleic acids of proliferating lymphocytes collected from different experimental mice. At day 7 post last immunization, splenocytes were stimulated with 1 μg/mL of OMVs. The assay was performed in triplicate, and data represent Mean stimulation index ± SE of two independent experiments. Asterisks indicate a statistically significant difference (\*\**P* ≤ 0.01) with respect to control mice (PBS group). **(B)** A dose-dependent increase of NO production in the culture supernatant of splenocytes treated with OMVs. One week post last immunization, splenocytes collected from mice belonging to different feeding groups were treated with varying concentrations of OMVs (0.1 μg/mL, 0.5 μg/mL, and 1 μg/mL) for 48 h. After treatment, culture supernatants were collected and assayed for NO production with the Griess reagent. The assay was performed in triplicate, and data represent Mean NO production ± SE of two independent experiments. Asterisks indicate a statistically significant difference (\**P* ≤ 0.05, \*\**P* ≤ 0.01) with respect to the control mice (PBS group).

### CS-OMVs immunization triggers NO production *in vitro*

Splenocytes harvested from experimental mice in response to *in vitro* stimulation with varying concentrations of purified OMVs results in high-level production of NO in the culture supernatants of cells collected from both CS-OMVs and IFA-OMVs immunized mice (Immunized Vs Control) (*P* ≤ 0.01). A critical analysis suggests a concentration-dependent (0.1 μg/mL, 0.5 μg/mL, 1 μg/mL) increase in NO production in response to *in vitro* stimulation with OMVs in immunized mice (Fig 4B).

### Immunophenotyping of OMVs specific T cell subsets (CD3^+^, CD4^+^, CD8^+^, and CD196^+^ T cells) in immunized mice

Immunophenotyping of T cell subsets (Th, Tc, and Th17 cells) in mice spleen collected from different experimental groups by flow cytometric analysis indicates a marked increase in total CD3^+^ T cells population in CS-OMVs (~32 %) and IFA-OMVs (~34 %) administered mice as compared to control animals (PBS: ~22 %; CS: ~25 %) (*P* ≤ 0.01) (Fig 5A, Table 3). With respect to Th and Tc cells, a significant rise in both CD4^+^ T (*P* ≤ 0.05) and CD8^+^ T (*P* ≤ 0.01) subset population was noted in the CS-OMVs group followed by IFA-OMVs immunized group (Immunized Vs Control). In contrast, a specific increase in CD196^+^ T cells population in IFA-OMVs immunized group was noted (*P* ≤ 0.01), while only a marginal rise was found in mice that received mucosal administration of CS-OMVs (Fig 5B, Table 3).

**Fig 5.**
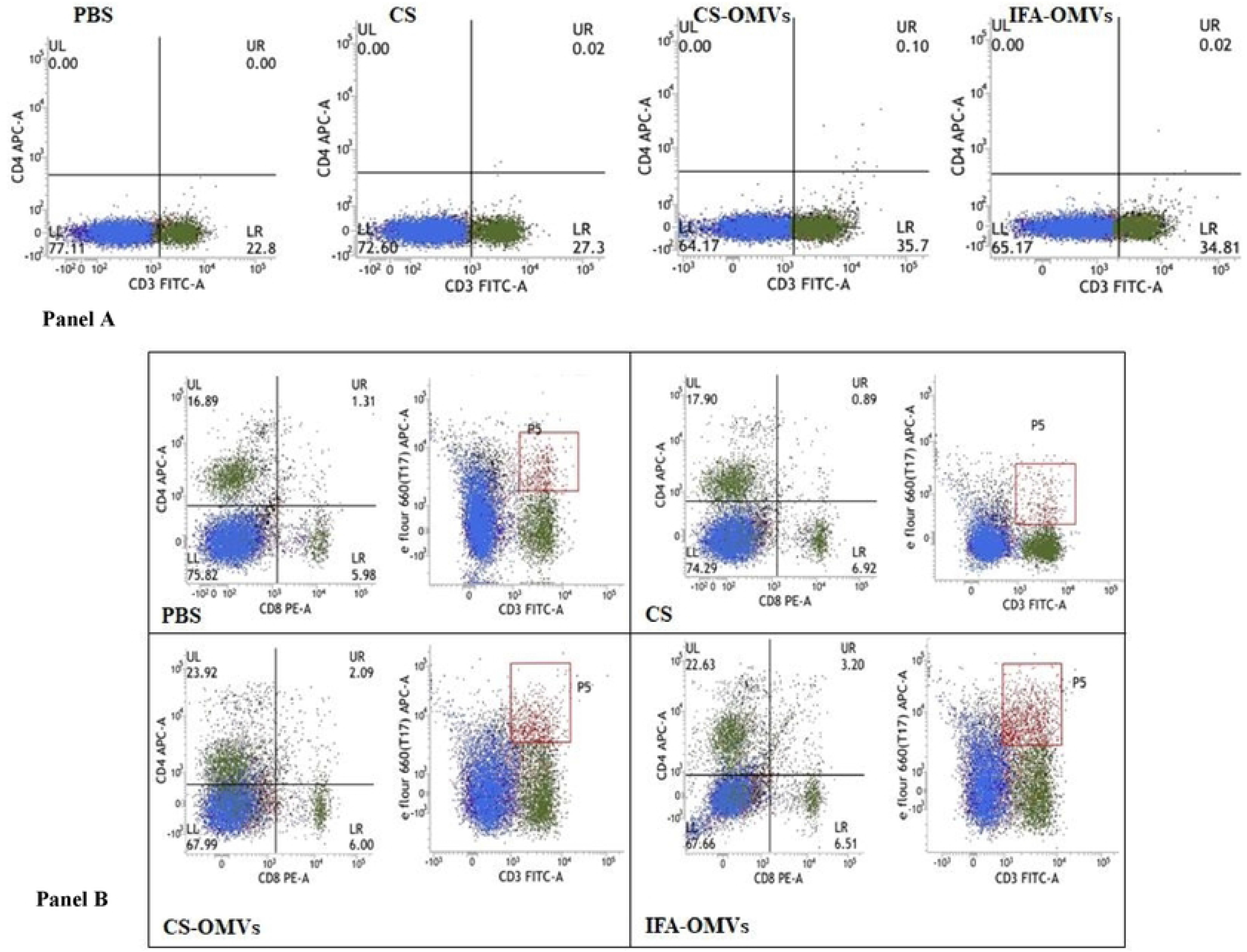
Immunophenotyping of T cells and their subsets by Flow cytometry. Splenocytes collected at day 7 post last immunization from mice belonging to immunized and unimmunized groups were stained with CD3-FITC, CD4-APC, CD8-PE, CD196-eFluor 660 monoclonal antibodies to count the population of total T cells, Th cells, Tc cells, and Th17 cells respectively. Lymphocytes were gated on the basis of their FSC-A and SSC-A. **(A)** Flowcytometric analysis showing a gated plot for total T cells (CD3 FITC) in splenocytes obtained from different experimental groups. For immunophenotypic profiles of Th and Tc cells, triple-staining was performed (CD3-FITC, CD4-APC, CD8-PE), whereas, for Th17 cells, double-staining was done (CD3-FITC, CD196-eFluor 660). (**B)** Gated plot for Th cells (CD4 APC), Tc cells (CD8 PE), and Th17 cells (eFlour 660) for various experimental groups. Channels FL1, FL2 were used as filters or detectors for FITC, PE-labeled antibodies, respectively, whereas channel FL4 was common for APC and eFlour 660 labeled antibodies.

### CS-OMVs immunization mediates pro-inflammatory cytokine responses in mice

Comparative analysis of mean fold changes of cytokine gene expression suggests transcriptional upregulation of IL-6 (*P* ≤ 0.05) and IFN-γ (*P* ≤ 0.01) genes in mice mucosally administered with CS-OMVs followed by IFA-OMVs injected mice (Immunized Vs Control). However, no changes were observed in IL-4 gene expression. Additionally, to see the effect of OMVs immunization of mice in modulating the innate immune response, TLR 4 gene was chosen; however, only minor changes were noted (Fig 6).

**Fig 6.**
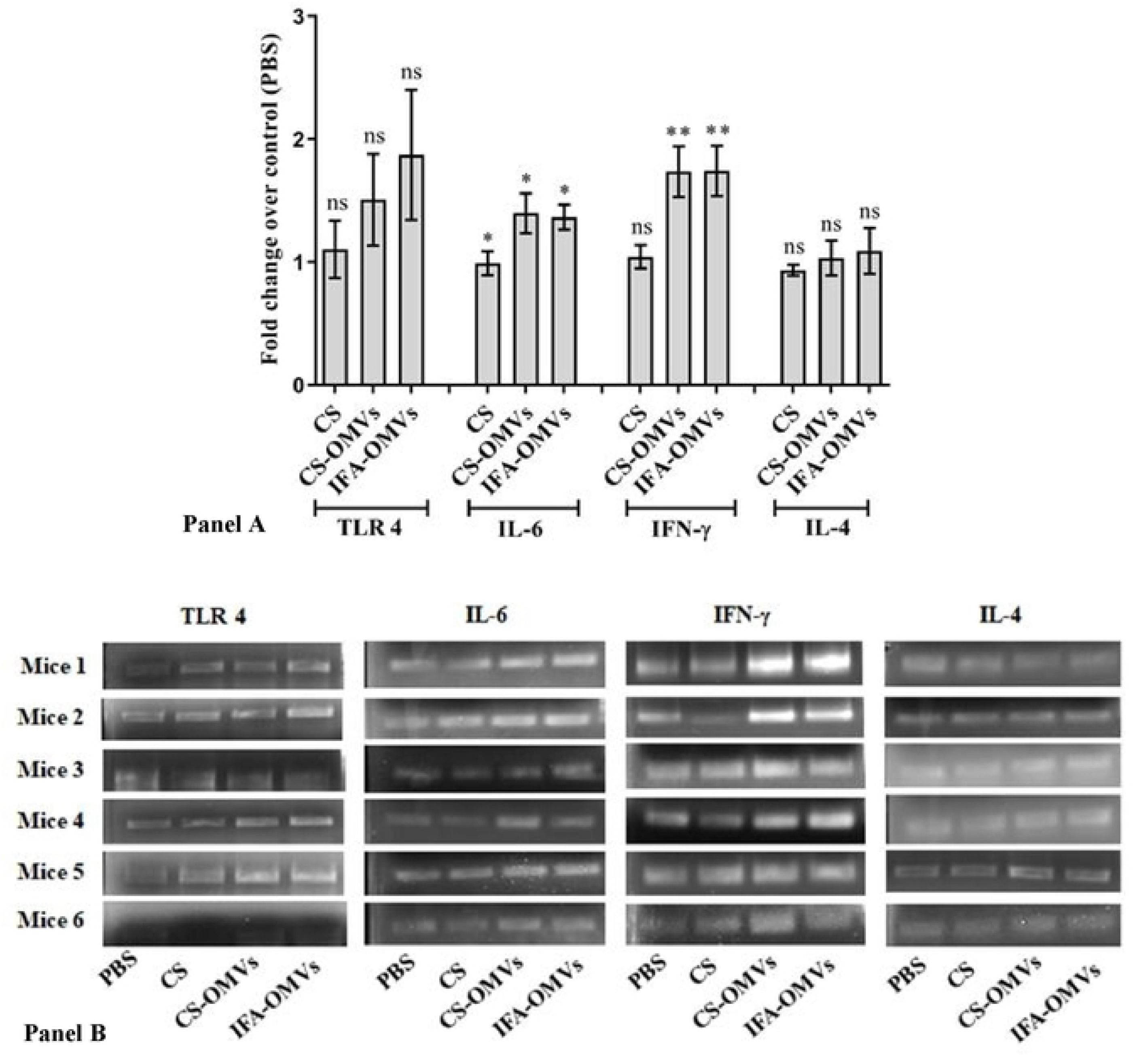
Comparative analysis of gene expression profile in experimental mice. Mesenteric lymph nodes collected at day 7 post last immunization were processed for transcriptional analysis of TLR 4, IL-6, IFN-γ, and IL-4 genes. **(A)** Fold changes of gene expression in response to immunization with only CS, CS-OMVs, and IFA-OMVs. Fold changes were calculated with respect to the control group (mice received PBS only). Data represent Mean fold change ± SE of two independent experiments. Asterisks indicate a statistically significant difference (\**P* ≤ 0.05, \*\**P* ≤ 0.01) with respect to the PBS group. (**B)** Agarose gel images (1 % agarose) of PCR amplified products using gene-specific primers showing the expression of mRNA of TLR-4 and cytokines (IL-6, IFN-γ, and IL-4). Data represent gel images of two independent experiments.

### Mucosal administration of CS-OMVs reduced cecal load of *C. jejuni* in challenged mice

To assess the protective efficacy of mucosal delivery of chitosan-coated OMVs, mice of different experimental groups were challenged with *C. jejuni,* and the bacterial load was determined in cecum at day 7 post-challenge. Although the data shows a similar trend with respect to total bacterial load (CFU/gm), we noted some variations between similar treatment groups within experimental repeats; hence, normalized data were used for comparative analysis (S3A Fig, Table S2). Critical analysis of combined and normalized data suggests a significant reduction in cecal load of *C. jejuni* in mice specifically immunized with CS-OMVs (~2 fold) followed by IFA-OMVs (~1.9 fold) injected mice (Immunized Vs Control) (*P* ≤ 0.01) (Fig 7A).

**Fig 7:**
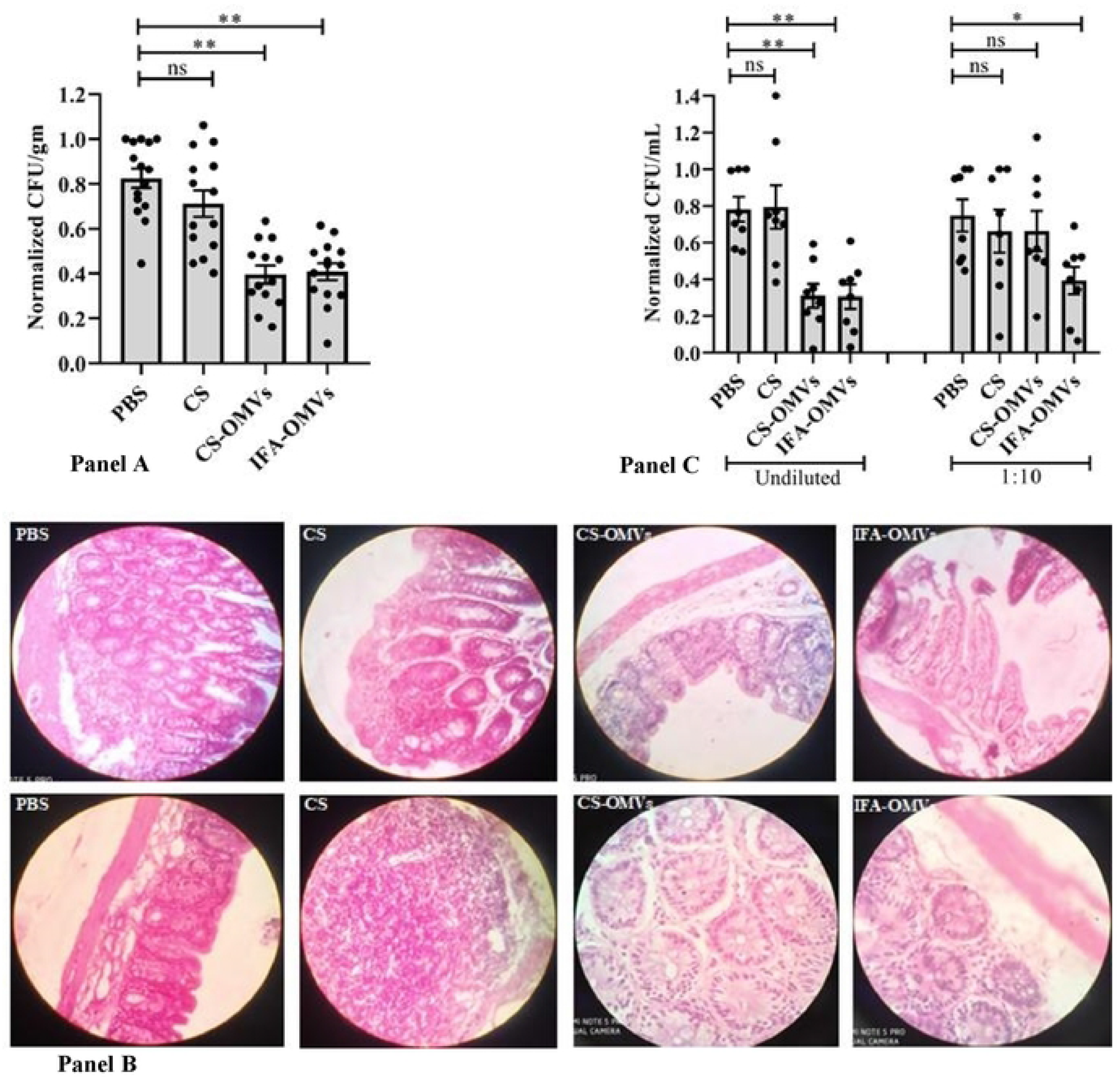
Immune-protective efficacy and histopathological changes in mice immunized with OMVs and challenged with *C. jejuni*. **(A)** Experimental mice were challenged with highly pathogenic 18aM *C. jejuni* isolate (1 × 10^8^ CFU/mice) and sacrificed at day 7 post last immunization. The normalized CFU/gm of cecum/mice in each experimental group was determined at day 7 post-challenge with *C. jejuni*. The number of viable *C. jejuni* recovered from the cecal content of different experimental groups showed a significant reduction in the bacterial load of mice orally administered with CS-OMVs followed by those received subcutaneous injection of IFA-OMVs as compared to the control groups (CS or PBS only). Data represent normalized CFU/gm ± SE of three independent experiments. Asterisks indicate a statistically significant difference in comparison to the PBS control group (\*\**P* ≤ 0.01). **(B)** Representative images of histopathological changes in cecal tissue collected at day 7 post-challenge with *C. jejuni.* Haematoxylin-and-Eosin (H&E) stained tissue sections showed desquamation of villi, focal necrosis, and degeneration of payer’s patches in mice administered with CS or PBS only. However, cecal tissues from mice immunized with CS-OMVs and IFA-OMVs indicates only minor changes in the morphology of villi and tissue architecture with diffuse infiltration of plasma cells. **(C)***In vitro* neutralization of *C. jejuni* by OMVs specific local antibody (sIgA) present in the faecal soup of mice immunized with OMVs either orally or systemically showed a significantly low number of *C. jejuni* (adhered + invaded) associated with human INT407 cells compared to the control groups. Data represent normalized CFU/mL ± SE of two independent experiments. Asterisks indicate a significant difference (*P ≤ 0.05, **P ≤ 0.01) statistically with respect to the PBS group.

### Mucosal administration of CS-OMVs prevents cecal pathology in challenged mice

The cecal tissues of challenged mice immunized with CS-OMVs or IFA-OMVs, except for some minor changes, no noticeable pathology was detected (Fig 7B). In contrast, unimmunized mice (PBS or CS group) challenged with *C. jejuni* showed marked inflammatory changes in cecal tissue, including lymphoid depletion, necrosis and degenerative changes mainly in the payer’s patches. Some desquamation of the villi with characteristics focal enterocytosis related pseudo-stratification was also clearly visible (Fig7B). Additionally, focal edema in sub-muscularis mucosae with some mononuclear and polynuclear cell infiltration was observed in unimmunized mice. Moreover, focal changes in terms of hyperplasia of goblet cells along with the accumulation of mucins and vacuolar degeneration in the cytoplasm of cells of the crypt, cryptal deformation were evident in mice that received CS only. Notably, diffuse plasma cell infiltration along the periphery of the crypt and accumulation of the proteinaceous material in the cells of the crypt was distinctly found in the tissue sections of immunized mice that received either CS-OMVs or IFA-OMVs (Fig7B).

### *In vitro* neutralization of *C. jejuni* by OMVs specific local (sIgA) antibody

Neutralization of *C. jejuni* with faecal soup (undiluted) collected from mice mucosally administered with CS-OMVs showed a significant reduction in the total number of associated bacteria (adhered + invaded) recovered from infected INT407 cells followed by IFA-OMVs immunized group (*P* ≤ 0.01) (Fig 7C, S3B Fig, Table S3).

## Discussion

Harmonized secretion of major virulence factors is a shared mechanism by many mucosal pathogens, including *C. jejuni*. However, except T6SS, C. *jejuni* lacks many classical virulence-associated secretion and export systems in comparison to the other enteric pathogens [10]. As an alternative means, *C. jejuni* employs OMVs as a concerted strategy to deliver active toxins and secretory proteins into the target cells [9]. With the ability to shuttle molecules between cells, OMVs are known to facilitate cell-to-cell communication and perhaps helps bacteria to limit their elimination during gut transit. With these unique functional diversities exhibited by OMVs, results of our *in vitro* invasion study suggest the possible involvement of naturally secreted OMVs in the enhancement of *C. jejuni* invasion of host cells in a dose-dependent manner and subsequent pathogenesis in a way similar to other enteric pathogens [39,40]. The ability of cell invasion by *C. jejuni* is likely to be facilitated by the presence of OMVs associated several serine proteases including high-temperature requirement protein (HtrA), Cj0511 as well as Cj1365c proteins, which are known to cleave E-cadherin adherence junction without affecting the fibronectin receptor of polarized as well as non-polarized cells lines including human IINT407 cells [12,41–43].

Given the significance of this system in overall defense, bacterial pathogenicity, and their ability to manipulate both B and T cell responses, the use of bacterial OMVs as possible tools for diverse biotechnological applications, including vaccine development against typical intracellular bacteria has raised significant attention in the recent times [44–52]. Because *C. jejuni* primarily adhere to intestinal epithelial cells, an upshot of vaccination against *C. jejuni* largely relies on the significant induction of local immune responses [24]. To this end, we described here a systematic approach to establish the value of CS coating over OMVs for promoting OMVs mediated *in vivo* immune-protection against *C. jejuni* challenge in the murine model [28–38].

The advantage of CS as naturally available mucoadhesive polymers for mucosal vaccine delivery platform has been conceptualized by several studies in the past [53]. Specifically, our group has recently demonstrated a comprehensive mucosal immunization modality using CS encapsulated haemolysin co-regulated protein (hcp) of T6SS in blocking cecal colonization of *C. jejuni* in chickens [54].

The outer membrane vesicles used in this study was isolated from a highly pathogenic *C. jejuni* (maintained in our lab) harbouring several genes encoding virulence factors (GEVFs), including *hcp* gene of *C. jejuni* T6SS. Since improper use of chemicals is often associated with the loss of lipoproteins and polysaccharides content, we purified OMVs without chemical treatment and confirmed their structural and morphological integrity [10,11,55,56]. As a polycationic polymer, CS is expected to form a positive coating around the negatively charged OMVs by electrostatic interaction which could be clearly evident from DLS data presented herein (Table 2) [57]. Expectedly, our biophysical analysis of CS coated OMVs suggest a marked increment in the mean hydrodynamic diameter of OMVs (149.9 to 221.9 nm) along with a significant drop in net negative charges (−23.4 to −8.2 mV). The negative charges of OMVs are primarily attributed to the LPS content of OMVs [15,58]. With our DLS data, additional verification of morphological features by SEM and TEM for both OMVs and CS-OMVs further substantiated the spherical nature of OMVs used in this study [15,40]. Taking into account the risk of premature degradation and intra-gastric instability of OMVs in the harsh gut environment, in addition to effective adsorption of local antigen-presenting cells (APCs), CS coating is expected to protect the OMVs from pH variability and enzymatic degradation. In fact, data presented here with respect to morphology and sizes support the enhanced stability of CS-OMVs over un-coated OMVs at various physiological conditions (Table S1) [59].

Since the LPS content of OMVs remains as one of the major concerns for OMVs based vaccine delivery platform, prior to *in vivo* study, we tested the cytotoxicity of OMVs in human INT407 cells and confirmed the non-toxic nature with the higher safety profile for OMVs (CT_50_ >100 μg/mL) (S2 Fig). Additionally, since no live bacteria were present during isolation and purification of OMVs from culture supernatant, OMVs based vaccine platform is expected to be safe.

Considering the non-invasive nature, simplicity of administration, and the importance of local immune responses against *C. jejuni,* we preferentially used the mucosal route for the present study. Significant increment of local (sIgA) and systemic (IgG) antibody responses in the intestines and serum collected from OMVs immunized mice either mucosally or systemically confirmed the *in vivo* immunogenicity of OMVs. High-level sIgA expression in the intestine could be credited to the successful interaction of CS-OMVs with the locally available APCs due to strong mucoadhesive property and high density of positive charge of CS [60].

As a non-complement fixing antibody isotype, the neutralizing ability of sIgA is in part, exhibited by binding of bacterial epitopes with glycans, generally associated with the secretory chain and constant region of α chain of sIgA, which in turn prevent bacterial adhesion to epithelial cells. A considerably low number of *C. jejuni* recovered from the cells incubated with antibody-treated bacteria strongly endorse the functionality of sIgA induced by oral administration of CS-OMVs. Hence, the neutralizing ability of local antibody against *C. jejuni* could be beneficial in terms of protecting intrinsically delicate monolayer of intestinal epithelial cells from inflammatory damage caused by *C. jejuni* adherence and invasion [61–69]. Considering that CD4^+^ T cells primarily mediate the help to local B cells in generating neutralizing antibodies, *in vitro* blocking of *C. jejuni* adherence to host cells has led us to explore the correlates of the type of immune responses elicited by mucosal administration of OMVs [70,71].

The consistent and steady rise of IgG in the serum with balanced antibody isotypic profiles in mice mucosally administered with OMVs indicate mixed Th1/Th2 type responses, which could possibly be due to the intrinsic property of the proteins associated with OMVs as well as the adjuvant effect of CS [72–74]. However, it is not clear how mucosal administration of OMVs affects the systemic antibody responses; one possible reason could be systemic migration of antigen primed local B cells present in the gut-associated lymphoid tissue (GALT) via lymphatic circulation [14,75–77]. Since pathogen-specific expansion and contraction of immune cells are the hallmarks of the effective activation of the immune system [78,79], we evaluated the priming effects of OMVs immunization by systematic analysis of cell proliferation, ability to produce nitric oxide (NO), expression of cytokine genes, and finally immunophenotypic profile of T cells [19,44,80–82].

The data obtained from *in vitro* incorporation of 5-bromodeoxyuridine (BrdU) into nucleic acids of proliferating splenocytes measured the rate of spleen cell proliferation [83]. The marked increase in lymphocyte proliferation with the present immunization regime (three doses) suggests T cells are effectively primed *in vivo* by oral immunization as immunological memory might have been induced within the specific subtypes of T cells [84,85]. The priming effect of OMVs was further substantiated by a dose-dependent increase in NO production in the culture supernatant of splenocytes treated with OMVs [86–89].

In parallel to these generic observations, our immunophenotyping data of T cell population within splenocytes showed a significant increase in total T cells (CD3^+^) [CS-OMVs: ~32 %], Th (CD4^+^) [CS-OMVs: ~21 %], Tc (CD8^+^) [CS-OMVs: ~7 %], with minor increase in Th17 (CD196^+^) [CS-OMVs: ~5%] subsets compared to control group (received PBS only). However, the elevation in the T cell population was in comparable range of a systemically administered group of mice. Activation of these cells following priming with OMVs is a clear indication of effective processing and presentation of protein antigens associated with OMVs when administered either locally or through a systemic route [90]. Given that CD4^+^ Th cells play a crucial role in vaccine-induced immunity, we submit that mucosal delivery of CS coated OMVs could facilitate differentiation of the naïve T cells into distinct functional subsets, including Th1. Moreover, considering the role of Th17 responses in immune-protection against several enteric pathogens, including *C. jejuni*, a modest increase in the population of Th17 subsets shows an additional advantage of OMVs immunization [91–94]. Together, immune phenotypic profiles of each T cell subset seem to be prudently tailored by the present immunization modality in regulating pathogen-specific immune responses [95,96]. Nevertheless, with the fact that once antigen has been cleared, memory T cells become the sole source for subsequent immune surveillance both locally and systemically, further study with a longer post-immunization timeline is required [97].

Critical analysis of the antibody isotype data obtained from mucosally administered mice while suggesting skewing of the Th cell differentiation towards mixed Th1/Th2 type responses, the transcriptional upregulation of IFN-γ, and IL-6 genes also supports the protective Th1 type response [98–100]. Intriguingly, we noted relatively low-level expression of IL-4 as Th2-inducing cytokines produced by the T cells. In spite of strong IL-6 expression, as a potent regulator for differentiation of naive CD4^+^ T cells to the Th2 phenotype, low-level expression of the IL-4 indicates that the IL-6 may trigger the Th2 pathway by inducing a small amount of endogenous IL-4. This could be presumably due to the autocrine differentiation factor for the Th 2 cells [101].

Considering that the LPS (or LOS) is the most abundant pathogen-associated molecular patterns (PAMPs) associated with OMVs [90], we next studied whether OMVs immunization has any role in TLR gene activation. However, only a minor increase in the TLR 4 gene expression was noted; this could be presumably due to less amount of LOS present in the OMVs. Nevertheless, purification methods, the time point for isolating OMVs, and differences among *C. jejuni* strain with respect to LOS content could be other determining factors for the final LOS content of OMVs used in this study [102,103].

Our final aim was embodied to see the effect of cellular and local immune responses towards immune-protection following challenge with highly pathogenic *C. jejuni*. The observed reduction in the cecal load of *C. jejuni* in mice belonging to either mucosally (~2 fold) or systemically immunized (~1.9 fold) groups was found to be significant compared to the controls. Further, a strong correlation between cecal load of *C. jejuni* with the degree of pathology in cecal tissue of immunized and unimmunized mice was observed, which suggests the ability of OMVs immunization in protecting delicate intestinal epithelium lining against *C. jejuni* invasion. Intriguingly, the infiltration of OMVs specific plasma cells in the intestinal follicles in immunized mice could be due to effective affinity maturation and class switching of antibody-producing B cells to IgA-secreting plasma cells [104–107].

Although no detectable difference in the magnitude of overall immune responses was observed in terms of route of administration of OMVs, considering the non-invasive nature, simplicity of administration with broad coverage of local and systemic immune responses, our approach of mucosal delivery of OMVs could be a safer alternative to the systemic mode of immunization. Notwithstanding that there are many factors that can influence the response to mucosal vaccination, the studies reported herein provide relevant insight of using chitosan to modulate OMVs specific immune-protection towards controlling the risk of systemic dissemination of *C. jejuni*.

## Materials and Methods

### Bacterial strains, culture conditions, cell lines, and other reagents

*C. jejuni* (18aM) was isolated from the cecal content of commercial broiler chickens and processed as per the procedure described elsewhere [54]. Briefly, samples were serially diluted in autoclaved distilled water and 0.1 mL from 10^−3^ dilution was plated onto Blood Free Campylobacter Selectivity Agar Base media (HiMedia, India) having CAT Selective Supplement (cefoperazone 8 mg/L, amphotericin 10 mg/L, and teicoplanin 4 mg/L) (HiMedia) followed by incubation for 48 h. Colonies found positive for *C. jejuni* were next grown in Mueller Hinton broth (HiMedia) supplemented with CAT supplement for 48 h under microaerophilic condition (10 % CO_2_, 5 % O_2,_ and 85 % N_2_) at 42 °C using microaerophilic generating gas pack (Anaerogas pack 3.5 L, HiMedia)

All chemical substances and reagents used in this study were of analytical grade. Chitosan (CS) and Sodium Tripolyphosphate (TPP) were obtained from HiMedia and Loba Chemie (India), respectively. Bicinchoninic acid (BCA) protein assay kit was purchased from Pierce Chemical Co. (USA). Incomplete Freund’s adjuvant and Histopaque 1077 were procured from Sigma (USA). Goat anti-mouse IgG (H+L; HRP) was purchased from Life Technologies (USA). HRP conjugated goat anti-mouse IgG1, IgG2a, IgG2b, and IgA antibodies were obtained from Bethyl Laboratories (USA). The FITC, APC, PE, eFlour 660 conjugated anti-mouse CD3, CD4, CD8a, and CD196, respectively, were obtained from eBioscience, Invitrogen (USA). Enzyme substrate 3, 3′,5, 5′-tetramethyl benzidine (TMB) was purchased from HiMedia. The human non-polarized INT407 cell line was procured from National Centre for Cell Science, Pune (India), and maintained in our laboratory following standard protocol. Primers used in the present study were purchased from IDT technologies (USA).

### Isolation and purification of OMVs derived from *C. jejuni* isolate

The vesicles were isolated from the culture supernatant of 18aM *C. jejuni* isolate following the method as described earlier with some modifications [108]. Briefly, cultures were grown for 14 h under the microaerophilic condition at 42 °C followed by centrifugation at 10,000 x g for 15 min at 4 °C. The supernatant was filtered through a 0.45 μM pore size membrane (Millipore) to remove the remaining bacterial cells. Approximately 0.1 mL of the filtrate was plated onto Mueller Hinton (MH) agar plate to test the presence of viable *C. jejuni* cells. In all cases, colonies were not observed. Vesicles recovered by ultracentrifugation of filtered supernatant at 150,000 x g for 2 h at 4 °C using a Ti 45 rotor (Beckman Instruments, USA), were washed with phosphate-buffered saline (PBS) and resuspended in PBS and stored at −20°C. The protein concentration of isolated OMVs was determined using the bicinchoninic acid (BCA) method.

### Role of OMVs in *C. jejuni* invasion of human INT407 cells

To assess the role of OMVs in *C. jejuni* invasion of the host cell, a gentamicin protection assay was performed according to the procedure mentioned previously in our lab [109]. Briefly, INT407 cells were seeded at a density of 3 x 10^5^ cells/well in a 24-well cell culture plate. A confluent monolayer of cells was treated with different concentration of OMVs (5 μg/mL, 10 μg/mL, 20 μg/mL) followed by co-incubation with *C. jejuni* (18aM isolate) at Multiplicity of Infection (MOI) 1:100 for 3 h at 37 °C and 5 % CO_2_ pressure. Post 3 h, the media was aspirated, followed by washing with PBS. For gentamicin protection assay, cells were treated with 150 μg/mL of gentamicin (prepared in 1X PBS) to kill the extracellular adhered bacteria and incubated for another 2 h at 37 °C under 5 % CO_2_ pressure. After incubation, *C. jejuni* infected monolayers were washed with PBS and lysed with 1 % Triton X-100 (prepared in PBS). The recovered bacteria were serially diluted in MH broth and plated onto MH agar plate followed by incubation at 42 °C for 24 h under the microaerophilic conditions in a tri-gas incubator (Thermo Scientific). Bacterial colonies that appeared on the plate were counted to enumerate colony-forming units (CFU). The assay was performed in triplicate and regression value was calculated using non-linear regression in GraphPad Prism software. Data represent Mean CFU/mL ± SE of two independent experiments.

### Immunogen preparation

#### Preparation of chitosan-coated OMVs for mucosal delivery

For mucosal delivery, chitosan was cross-linked with sodium tripolyphosphate (CS-TPP) following the protocol described previously in our lab with some modifications [54]. Briefly, 700 μL of freshly isolated OMVs (~70μg) was mixed with 3.3 mL of CS solution (0.05 % w/v) followed by drop-wise addition of 1 mL of TPP solution (0.1% w/v). The mixture was stirred for 2 h at 4 °C. The slurry formed was centrifuged at 14,000 x g for 30 min and resuspended in 0.2 mL of sterile PBS (pH 7.4).

### Biophysical characterization of CS coated OMVs

#### Dynamic light scattering (DLS) and measurement of zeta potential (ζ)

The size distribution and overall surface charge (zeta potential; ζ) of OMVs and CS coated OMVs were measured as described elsewhere with minor changes [110,111]. Concisely, OMVs alone and CS coated OMVs were diluted (1:100 dilution) in PBS for size and milliQ water for charge analysis followed by sonication for 15 min. After sonication, samples were analyzed for size distribution by DLS and zeta potential by laser doppler micro-electrophoresis using Malvern Zetasizer Nano ZS instrument (USA). In addition, to confirm the stability of NPs coated OMVs, size distribution analysis was performed at varying pH, buffer compositions, and incubation time points (details of the experiment are mentioned in Table S1).

#### Scanning electron microscopy (SEM)

The morphology of the isolated OMVs and CS-OMVs were examined through SEM image analysis (Carl Zeiss SUPRA 55 V P FESEM). Samples were processed according to the method mentioned previously with some modifications [112]. Briefly, for OMVs, specimens were fixed overnight in 2.5 % (v/v) glutaraldehyde (prepared in PBS; pH 7.4) at room temperature (RT). Fixed samples were washed thrice with PBS for 10 min each followed by sequential dehydration in 35 %, 50 %, 70 %, 95 % ethanol for 10 min each and 100 % ethanol for 1 h for complete dehydration. Finally, fixed and dehydrated samples were vacuum-dried overnight. For CS-OMVs, NPs were first dispersed in milliQ water (1:100 dilution) and sonicated for 15 min in a bath sonicator (Thermo Fisher Scientific, USA). Dispersed samples were further processed following the procedure as mentioned above for OMVs. Samples were thoroughly dried under vacuum and fixed to aluminum stubs with silver conductive paint and sputter-coated with gold and examined using a Supra 55 Carl Zeiss scanning electron microscope. Images were analyzed using ImageJ software.

#### Transmission electron microscopy (TEM)

For the TEM analysis, OMVs and CS-OMVs samples were diluted in milliQ water (1:100 dilution) followed by sonication for 30 min at RT. After 30 min, ~ 5 μL of the sonicated samples were drop cast onto 300-mesh formvar carbon-coated copper grids (Electron Microscopy Sciences, UK) and negatively stained with UranyLess (Electron Microscopy Sciences). Samples were left undisturbed for 10 min followed by removal of excess fluid using filter paper. The samples were vacuum-dried overnight. Data acquisition was made using a JEOL JEM-2100 Plus LaB6 series electron microscope (Japan) operating at an accelerating voltage of 120 kV. Images were analyzed using ImageJ software.

### Assessing the immune-protective potential of OMVs in mice

#### Immunization and challenge schedule

Female BALB/c mice were purchased from the National Centre for Laboratory Animal Sciences, National Institute of Nutrition, Hyderabad, India. Six-week-old mice (20±1 g) were separated into randomized groups of 10 animals (n=10 per group). Mice were divided into four experimental groups as follows: Group 1: PBS (Sham control); Group 2: CS (Vehicle control); Group 3: chitosan-coated OMVs (CS-OMVs); Group 4: OMVs emulsified with Incomplete Freund’s adjuvant (IFA-OMVs). Mice of experimental Groups 1 to 3 were immunized orally, whereas mice belonging to Group 4 were injected through a subcutaneous route. Group 3 and 4 received 20 μg of OMVs in 100 μL PBS for all immunizations. At day 7 post last immunization half of the animals belonging to different groups were sacrificed for sample collection, whereas the remaining half were challenged with 1×10^8^ CFU/mice of *C. jejuni*.

#### Sample collection

##### Blood

Approximately 80 μL blood samples were collected from mice by retro-orbital puncture with heparinized capillaries at day 7 post last immunization. The collected blood was allowed to clot at room temperature (RT) for 2 h followed by centrifugation at 1000 x *g* for 15 min. The separated sera were stored at −20 °C until use.

##### Faeces

Faecal pellets were obtained from individual mice at day 7 post last immunization and resuspended in IgA extraction buffer (1X PBS pH 7.4 containing 0.05 % Tween 20, 0.5 % fetal bovine serum, 1mg/ml EDTA, and 1 mM phenylmethanesulfonyl fluoride (PMSF) as protease inhibitor). Pellets were vortexed until thoroughly macerated, and then the insoluble material was pelleted by centrifugation at 1000 x g for 20 min at 4 °C. The clarified faecal extracts were stored at −20 °C until further use.

##### Intestinal lavages

The lavage was collected at day 7 post last immunization and processed as described previously with slight modifications [113]. Briefly, the ileocecal junction was cut off from each sacrificed mice, and the interior of the intestine was flushed with 0.2 mL PBS. After centrifugation at 1000 × g for 15 min at 4 °C, the supernatants were collected and stored at −20 °C until use.

##### Tissue samples

Spleen and mesenteric lymph node (mLN) were collected from 5 mice of each experimental group at day 7 post last immunization under sterile condition. Spleen samples were immediately processed for cell-mediated immune response study, whereas mLN were stored in RNA later (Qiagen, USA) at − 20 °C till further use.

### Local and systemic antibody responses in mice immunized with OMVs

#### Mucosal antibody responses (sIgA) in gastric lavages and faecal soups

The production of OMVs specific secretory IgA (sIgA) antibody was measured in intestinal lavage and faecal soup of the immunized mice by indirect ELISA as mentioned elsewhere with minor changes [114]. In brief, 96-well ELISA plates (Thermo Fisher Scientific) were coated with OMVs (100 ng/well) overnight at 4 °C followed by washing with 1X PBS-T (0.05% Tween- 20 in PBS) and blocking with 3 % Bovine Serum Albumin (BSA) (prepared in PBS-T) at 37 °C for 1 h. Next, wells were washed with PBS-T and incubated with two-fold serially diluted lavage and faecal soup collected from different experimental mice (starting with 1:2 dilution) for 2 h at RT followed by another 1 h incubation with HRP conjugated Goat anti-mouse IgA secondary antibody (1:3000 dilution). Following several washes, the TMB substrate was added to each well. Finally, the reaction was stopped with 50 μL of 1 M H_2_SO_4,_ and the absorbance was read at 495 nm in a microplate reader (BioTek, USA). Data represent Mean of absorbance ± SE of two independent experiments.

#### Systemic antibody responses in sera

Serum antibody against OMVs was determined through indirect ELISA as mentioned in the above section except sera samples were two-fold serially diluted (starting from 1:20) followed by incubation with HRP conjugated Goat anti-mouse IgG, IgG1, IgG2a, and IgG2b secondary antibodies (1:3000 dilution). Subsequently, wells were treated with TMB substrate and finally, the reaction was stopped with 1 M H_2_SO_4,_ and the absorbance was read at 495 nm in a microplate reader (BioTek, USA). Data represent Mean of absorbance ± SE of two independent experiments.

### Cellular immune responses in mice immunized with OMVs

#### Preparation of mononuclear cells

To isolate mononuclear cells, mice from each experimental group were sacrificed, spleens were removed and lymphocyte enriched mononuclear cells were separated using Histopaque 1077 as mentioned elsewhere with slight modifications [115]. Briefly, spleens were collected under aseptic condition, washed thrice with PBS, and resuspended in 1 mL complete RPMI 1640 growth media (10 % fetal bovine serum and 1X penicillin-streptomycin). Next, tissue was transferred to a sterile petri-dish and crushed with a flat end of 5 mL disposable syringe. Cell suspension from the petri-dish was aspirated and filtered through a 70 μm cell strainer. The flow-through comprising single-cell suspensions was layered onto pre-warmed histopaque in a 1:1 ratio and centrifuged at 400 x g for 20 min at 23 °C. The middle cloudy whitish interface was taken off (containing mononuclear cells) and resuspended in complete RPMI growth media for further use in a splenocyte proliferation assay.

#### Splenocytes proliferation assay

To determine the splenocyte proliferation of experimental animals, spleen lymphocyte proliferation assay was performed using the BrdU cell proliferation ELISA kit following the manufacturer’s protocol (Abcam). Briefly, the lymphocyte-enriched mononuclear cells obtained as described in the above section were counted and seeded at a density of 2 × 10^5^ cells/ well in triplicate in flat-bottomed 96-well cell culture plates, co-stimulated with OMVs (1 μg/mL) and incubated at 37 °C under 5 % CO_2_ pressure. Splenocytes stimulated with mitogen Concanavalin A (ConA) (10 μg/mL), or RPMI 1640 media alone (un-stimulated) were kept as positive and negative controls respectively. Post 24 h of stimulation, 20 μL of 1X BrdU reagent (Abcam, USA) was added to each well and incubated for another 24 h. After incubation, cells were fixed and DNA was denatured using a fixing solution followed by probing with anti-BrdU monoclonal detector antibody for 1 h at RT. Next, cells were labelled with 1X HRP conjugated goat anti-mouse IgG antibody. After washing with wash buffer, the plates were developed with the TMB substrate. The reaction was stopped with stop solution and absorbance was read at 450 nm using Epoch 2 microplate reader (BioTek, USA). Data are expressed as stimulation index (S.I) which is described as the ratio of the mean absorbance of stimulated cells to that of unstimulated cells. The assay was performed in triplicate and data represent Mean stimulation index ± SE of two independent experiments.

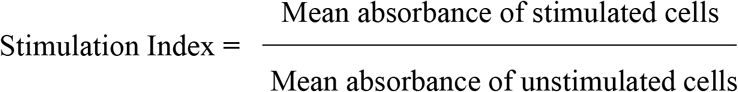

#### Assessment of Nitric Oxide (NO) production

To determine NO production in the splenocytes of experimental mice, the accumulation of nitrite was quantified using the standard Griess assay as described previously with minor modification [116]. Briefly, splenocytes were seeded at a density of 2 × 10^4^ cells/well in phenol red-free complete RPMI 1640 growth media in 12-well cell culture plate followed by stimulation with varying concentration of OMVs (0.1 μg/mL, 0.5 μg/mL, and 1.0 μg/mL). Cells with RPMI alone and cells stimulated with ConA (10 μg/mL) were kept as controls. After 48 h of incubation, 100 μL of media were incubated with an equal volume of Griess reagent (1 % sulfanilamide, 0.1 % naphthyl ethylenediamine dihydrochloride, 2.5 % H_3_PO_4_) for 15 min at RT. The absorbance was measured at 570 nm in Epoch 2 microplate reader (BioTek, USA). The conversion of absorbance into micromolar concentrations of NO was deduced from a standard curve using a known concentration of NaNO_2_. The assay was performed in triplicate, and data represent Mean NO concentration ± SE of two independent experiments.

#### Immunophenotyping of T cell population in immunized mice

To determine the total T cell and their subsets, mononuclear cells were isolated from the spleens as mentioned previously and processed for flow cytometry as described elsewhere with minor modifications [117]. Briefly, 1×10^6^ mononuclear cells were resuspended in 0.1 mL of FACS buffer (PBS with 1 % BSA) in flow cytometry tubes. Cell surface marker staining was performed by probing them with the following monoclonal antibody (mAb) combination: CD3-FITC (0.0025 μg/μL), CD4-APC (0.00125 μg/μL), CD8-PE (0.0025 μg/μL), CD196-eFluor 660 (0.0015 μg/μL) followed by 45 min incubation on ice in the dark. The proportion of T cell subsets in the spleen were specifically analyzed by selective gating based on the size and granularity of the cells using the FACSCalibur flow cytometer (BD Biosciences) and analyzed with the CellQuest Pro software. Data represent Mean cell percentage ± SE of two independent experiments.

#### Toll-like receptor 4 (TLR-4) and cytokine genes expression

The expression of cytokine genes was determined in RNA extracted from mLN tissue stored in RNA *later* as per the method described elsewhere with some modifications [118,119]. Briefly, 30 mg of the mLN tissue was washed with PBS and homogenized in Trizol reagent followed by 15 min incubation at RT. Next, chloroform was added in a 1:1 ratio and mixed vigorously (without vortex) and incubated for 15 min at RT to form layers. After incubation, the samples were centrifuged at 10,000 x g for 20 min at 4 °C. The upper aqueous layer was collected in a centrifuge tube followed by the addition of 500 μL of isopropanol, mixed and incubated at −20 °C overnight. The following day samples were centrifuged at 10,000 x g for 30 min at 4 °C. The pellet obtained was washed with 70 % ethanol and air-dried. Finally, the pellet was dissolved in nuclease-free water (NFW). The concentration of RNA was determined using Epoch 2 microplate spectrophotometer (BioTek).

Approximately 2 μg of RNA was used for cDNA synthesis using the Superscript Reverse Transcriptase kit following the manufacturer’s protocol (BioBharati, India). Primers used to assess the expression of genes are listed in Table 1. PCR amplification was carried out in a total volume of 20 μL master mix containing forward and reverse primers, dNTPs, Taq buffer, Taq polymerase, cDNA, and NFW. The PCR cycle consisted of initial denaturation at 94 °C for 3 min followed by 30 cycles of amplification at 94 °C for 1 min, 50 °C to 60 °C for 45 sec, 72 °C for 2 min followed by a final extension at 72 °C for 5 min. The expression of β-actin as a housekeeping gene was used for normalization of the data between samples. Data are presented as the mean fold change calculated with respect to the control group (received PBS only) using Image Lab™ software (Bio-Rad, USA). Data represent Mean fold changes ± SE of two independent experiments.

**Table 1.**
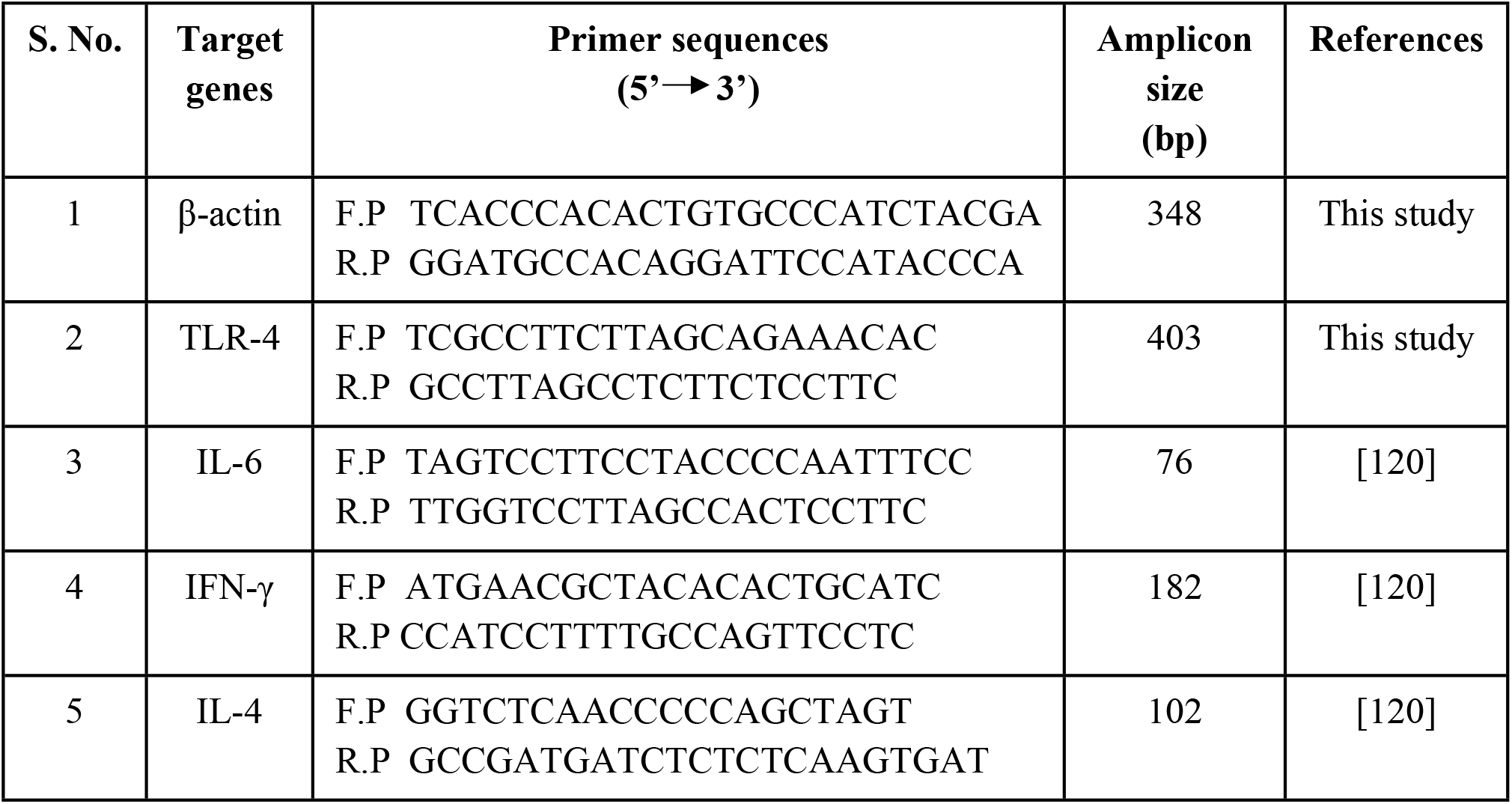
List of the primers used in the present study.

**Table 2.** Biophysical characteristics of OMVs and CS-OMVs by DLS, SEM, and TEM analysis.

| Types | Average size (nm) |  |  | Zeta Potential (ζ)<br>(mV) |
| --- | --- | --- | --- | --- |
|  | DLS | SEM | TEM |  |
| OMVs | 149.9 | 110 | 130 | -23.4 |
| CS-OMVs | 221.9 | 157 | 165 | -8.2 |

**Table 3.**
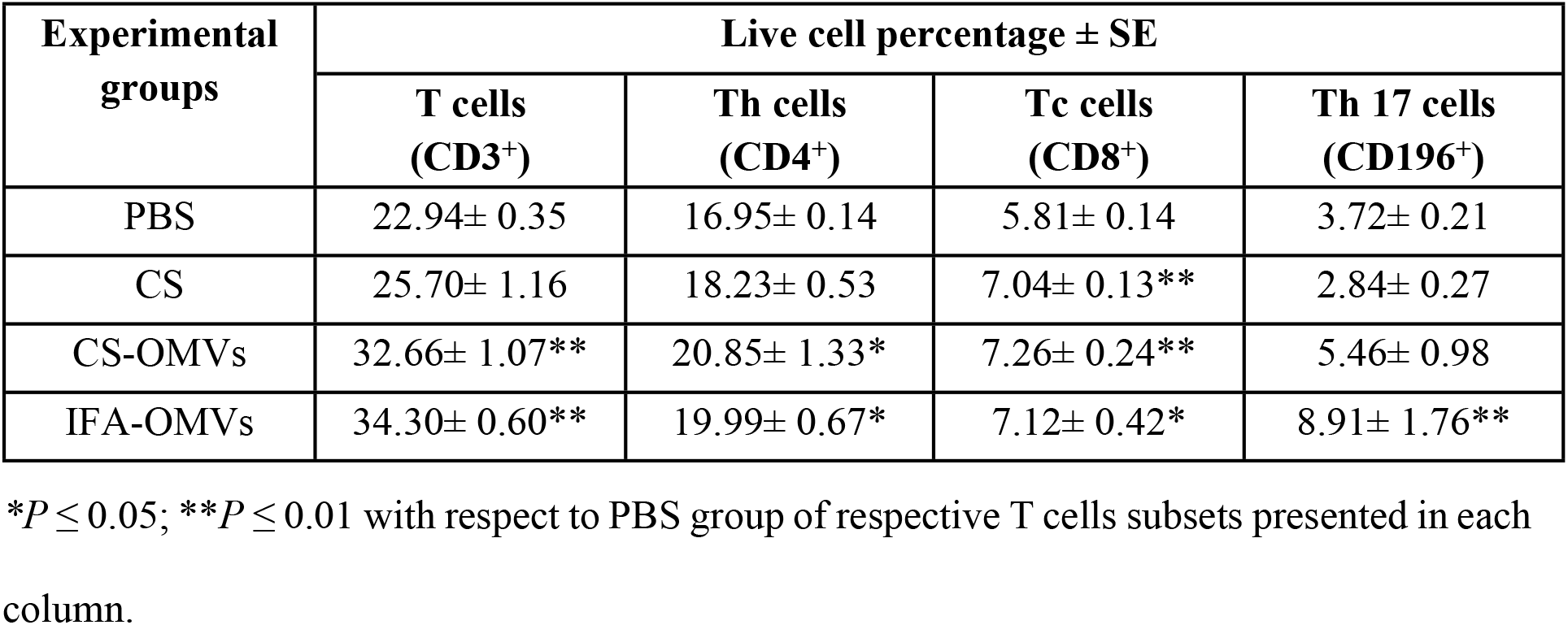
Mean percentage of total T cells and other subsets population (Th, Tc, and Th17) among different experimental groups.

| Experimental groups | Live cell percentage ± SE | | | |
| --- | --- | --- | --- | --- |
|  | T cells (CD3<sup>+</sup>) | Th cells (CD4<sup>+</sup>) | Tc cells (CD8<sup>+</sup>) | Th 17 cells (CD196<sup>+</sup>) |
| PBS | 22.94± 0.35 | 16.95± 0.14 | 5.81± 0.14 | 3.72± 0.21 |
| CS | 25.70± 1.16 | 18.23± 0.53 | 7.04± 0.13** | 2.84± 0.27 |
| CS-OMVs | 32.66± 1.07** | 20.85± 1.33* | 7.26± 0.24** | 5.46± 0.98 |
| IFA-OMVs | 34.30± 0.60** | 19.99± 0.67* | 7.12± 0.42* | 8.91± 1.76** |

\* $P \leq 0.05$ ; \*\* $P \leq 0.01$ with respect to PBS group of respective T cells subsets presented in each column.

#### Effect of OMVs immunization in cecal colonization of *C. jejuni*

One week post last immunization, mice of various experimental groups were challenged with 18aM *C. jejuni* isolate (1 × 10^8^ CFU/mice) in 100 μL of PBS. At day 7 post-challenge, mice from each group were sacrificed, the cecum was removed and cecal contents were homogenized in 2 mL of PBS (pH 7.4). Serial dilution of homogenized cecal content was made in MH broth, and 0.1 mL from 10^−3^ dilution was plated onto blood-free *Campylobacter* Selective Agar Base media plate having CAT supplement. All the plates were incubated at 42 °C under the microaerophilic conditions for 48 h. The number of colonies that appeared on the plate were expressed as normalized CFU/gm of the cecum ± SE of three independent experiments. The formula used for the normalization (N) of data is as follows:

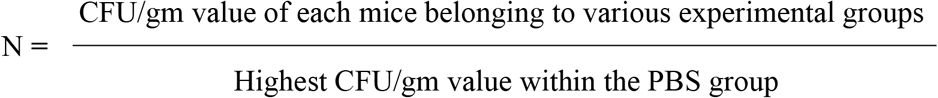

#### Histopathological analysis of cecal tissues

To determine histological changes, histopathological analysis of ceca from experimental mice at day 7 post bacterial challenge was performed as per the method described elsewhere with slight modifications [121]. Briefly, the cecum was fixed in 10 % formal solution (prepared in PBS) for 48 h at RT. The portion of the fixed specimens was washed overnight under running tap water. Following washing, samples were dehydrated sequentially using 70 %, 90 %, and 100 % ethanol for 1 h each for complete dehydration. Fixed and dehydrated tissue samples were next treated with clearing agents, xylene I and II, for 1 h each. Samples were further impregnated in paraffin I, paraffin II, and paraffin III each for 1 h. Finally, samples were embedded and sectioned at 4 μm. The samples were mounted on slides and stained with hematoxylin-eosin.

#### Assessment of the neutralization effect of secretory IgA on adherence and invasion of *C. jejuni* to INT407 cells

To determine the immune-protective efficacy of mucosal sIgA in blocking *C. jejuni* cell adherence and invasion, *in vitro* neutralization assay was performed as per the method described elsewhere with minor modifications [122]. Briefly, 1.5 × 10^6^ *C. jejuni* cells (18aM isolate) were co-incubated with undiluted and 10-fold serially diluted faecal soup for 2 h at 42 °C under the microaerophilic conditions in a tri-gas incubator. After incubation, human INT407 cells seeded at a density of 1.5×10^4^ cells/well in a 96-well cell culture plate were incubated with treated *C. jejuni* at MOI 1:100 for 3 h in 5 % CO_2_. After 3 h, cells were thoroughly washed with PBS and lysed with 1 % Triton X 100 (prepared in PBS). Lysed cells were serially diluted in MH broth and 0.1 mL from 10^−3^ dilution were plated onto MH agar plate followed by incubation at 42 °C for 24 h under the microaerophilic conditions. Colonies observed on the plates were counted for each experimental group to calculate CFU. Data represent normalized CFU/mL ± SE of two independent experiments.

#### Statistical analysis

The GraphPad Prism statistical software (Version 8) was used for graphical presentations and data analysis. The diameter of SEM and TEM images were examined using image J software. The regression (R^2^) value for invasion assay was calculated using a non-linear regression curve. Shapiro-Wilk test was performed to confirm the normal distribution. The Student *t*-test (two-tailed, unpaired) or non-parametric Mann-Whitney U test were performed to compare significance among various experimental groups. The \**P* ≤ 0.05, \*\**P* ≤ 0.01 were considered statistically significant.

#### Ethics statement

The mice experimentation protocol was approved by Institute Animal Ethics Committee (IAEC), Indian Institute of Science Education and Research Kolkata, and all procedures were conducted in accordance with the Committee for the Purpose of Control and Supervision of Experiments on Animals (CPCSEA) guidelines, MoEF & CC, Govt. of India. The permit number of the experimental protocols approved by the IAEC was IISERK/IAEC/AP/2020/50.

## Author contributions

AS and AIM designed the experiment, performed the experiment, analyzed the data, and wrote the manuscript. AK performed the biophysical characterization of OMVs and contributed to statistical analysis. TG assisted in flow cytometry data analysis and interpretation. SM contributed in the analysis and interpretation of histopathology data.

## Acknowledgments

The authors AS and AK thanks IISER Kolkata and UGC for providing fellowships, respectively. We acknowledge the central Electron Microscopy Facility at the IISER Kolkata for SEM and TEM images. We also thank Dr. Dipjyoti Das, Assistant Professor, Department of Biological Sciences, IISER Kolkata, for his input in the statistical analysis. We would also like to thank the Animal facility staff at the IISER Kolkata for assistance in animal experimentation.

